# Accurate identification of invasive *Aedes* mosquito species using low-cost imaging and geometric wing morphometrics

**DOI:** 10.64898/2026.04.08.717229

**Authors:** Felix Gregor Sauer, Hanna Jöst, Tatiana Sulesco, Pride Duve, Do Huy Loc, Paula Meyer, Kristopher Nolte, Renke Lühken

**Affiliations:** Bernhard Nocht Institute for Tropical Medicine, Hamburg, Germany; Kommunale Aktionsgemeinschaft zur Bekämpfung der Schnakenplage (KABS), Speyer, Germany; Institute of Tropical Medicine, University of Tübingen, and German Center for Infection Research (DZIF), Tübingen, Germany

## Abstract

Accurate species identification is crucial to assess the medical and veterinary relevance of a mosquito specimen, but it requires high experience of the observers and well-equipped laboratories. This study aimed to evaluate whether low-cost imaging in combination with geometric wing morphometrics can provide accurate identification of invasive, morphologically similar *Aedes* species.

The right wings of 670 female specimens covering 184 *Ae. aegypti*, 156 *Ae. albopictus*, 166 *Ae. j. japonicus* and 164 *Ae. koreicus*, were removed, mounted and photographed with a professional stereomicroscope (Olympus SZ61, Olympus, Tokyo, Japan) and a macro lens (Apexel-24XMH, Apexel, Shenzhen, China) attached to a smartphone. The coordinates of 18 landmarks on the vein crosses were digitalized by a single observer for each image. In addition, the landmarks of 20 specimens per species and imaging device were digitalized by six different observers to assess the degree of the observer error. The superimposed shape variables were used to compare the species classification accuracy of linear discriminant analysis (LDA), support vector machine (SVM), Random Forest (RF), and XGBoost.

In the single-observer landmark data, the LDA achieved the best classification results with a mean accuracy of 95 % for landmarks from microscope images and 92 % for those obtained from smartphone images. In the multi-observer landmark data, LDA consistently performed worse than the other three classifiers, and the reduction in the accuracy was more pronounced for smartphone images than for microscope images. This pattern was associated with a higher degree of observer error for smartphone images, as confirmed by a landmark-wise comparison across all landmarks.

Geometric wing morphometrics provides a reliable method to distinguish the most common invasive *Aedes* species in Europe. Thereby, the image quality obtained by smartphones equipped with a macro lens is sufficient and represents a cost-effective alternative to professional microscopes. However, due to the greater degree of observer variation for smartphone images, landmark coordinates for such images should ideally be collected by a single observer.

## Introduction

Since 2007, the number of *Aedes*-borne disease outbreaks has increased substantially in Europe (Cattaneo et al. 2025). This rise in autochthonous infections is closely linked to the spread of invasive *Aedes* species. The most prominent species in Europe is *Aedes albopictus*, which is rapidly spreading and has been responsible for several outbreaks of chikungunya virus, dengue virus, and Zika virus in Southern Europe (Lühken et al. 2023). In addition, *Ae. aegypti*, the primary vector of yellow fever virus and dengue virus in tropical regions, has been reported in Madeira (Seixas et al. 2019), Cyprus (Vasquez et al. 2023) and for areas surrounding the Black Sea (Yunicheva et al. 2008). Other invasive *Aedes* species in Europe include *Ae. japonicus japonicus* (from now *Ae. japonicus*) and *Ae. koreicus. Aedes japonicus* was first detected in Belgium in 2002 (Versteirt et al. 2009) and has since spread extensively, becoming established across large parts of Central and Southern Europe (Koban et al. 2019). In contrast, the closely related *Ae. koreicus* exhibits a more localized distribution, with small established populations in Belgium, Germany, Hungary and Italy (Kurucz et al. 2022). Compared with *Ae. albopictus, Ae. japonicus* and *Ae. koreicus* are likely to play a minor role for pathogen transmission in Europe. However, the vector competence of *Ae. japonicus* and *Ae. koreicus* has been experimentally confirmed for several pathogens, including Sindbis virus, western equine encephalitis virus, dengue virus, chikungunya virus, and *Dirofilaria immitis* (Schaffner et al. 2011, Montarsi et al. 2015, Jansen et al. 2021, Jansen et al. 2025).

*Aedes aegypti, Ae. albopictus, Ae. japonicus*, and *Ae. koreicus* produce drought-resistant eggs that are laid in small water bodies, such as tree holes, as well as in artificial containers, including used tyres or small buckets (Becker et al. 2020). These eggs can survive long dry periods and be transported globally, for instance through the international trade of used tyres (Hawley et al. 1987, Versteirt et al. 2009). At the same time, climate change is predicted to further expand their potential geographical distribution (Kraemer et al. 2019, Liu-Helmersson et al. 2019). Consequently, these four *Aedes* species have a high potential to spread into previously uninhabited European regions, underscoring the need for rapid detection and response measures to prevent population establishment. Therefore, accurate species identification is key to quickly detect local populations of invasive *Aedes* species. However, traditional morphological identification based on taxonomic keys requires extensive entomological expertise and is often compromised when specimens are damaged (Jourdain et al. 2018). This is particularly true for *Ae. aegypti, Ae. albopictus, Ae. japonicus*, and *Ae. koreicus*, which closely resemble each other and require well-preserved specimens for an accurate morphological species identification (Patsoula et al. 2006, Pfitzner et al. 2018). Alternative molecular identification methods require well-equipped laboratory facilities, limiting their applicability in resource-constrained settings.

Landmark-based geometric wing morphometrics has emerged as a promising alternative for mosquito species discrimination, including among closely related and morphologically similar *Aedes* species (Henry et al. 2010, Martinet et al. 2021, Sauer et al. 2023). The anatomical junctions of the mosquito wing veins are ideal targets for the collection of homologous landmark coordinates used in geometric morphometric analyses (Lorenz et al. 2017). However, although to a lesser extent compared to molecular techniques, the application of geometric wing morphometrics is also constrained by the need for specialized and often expensive equipment, such as stereomicroscopes and high-resolution cameras, to obtain sufficient image quality for clear visualization of the wing veins. Moreover, its broader implementation is challenged by observer-dependent variation in landmark digitization. This user effect can bias the results obtained from geometric morphometrics, thereby reducing reproducibility and hindering the transferability of species classification results across studies (Dujardin et al. 2010).

The aim of this study was to determine whether low-cost imaging can provide reliable and accurate species identification by means of geometric wing morphometrics. We applied landmark-based geometric morphometric methods to distinguish *Ae. aegypti, Ae. albopictus, Ae. japonicus*, and *Ae. koreicus*. Thereby, we compared the use of a professional stereomicroscope and a macro lens attached to a smartphone as a potentially cost-effective alternative for image collection. The resulting species classification accuracy was evaluated for four different classification algorithms. In addition, we quantified the degree of observer bias in the landmark collection associated with the different imaging devices.

## Methods

### Data set and image collection

The right wing of 184 *Ae. aegypti*, 156 *Ae. albopictus*, 166 *Ae. japonicus*, and 164 *Ae. koreicus* female specimens were removed and mounted with Euparal (Carl Roth, Karlsruhe, Germany). Subsequently, each wing was photographed with a camera (Olympus DP23, Olympus, Tokyo, Japan) connected to a stereomicroscope (Olympus SZ61, Olympus, Tokyo, Japan), and a smartphone (iPhone SE 3rd Generation, Apple Inc., Cupertino, CA, USA) equipped with a macro lens (Apexel-24XMH, Apexel, Shenzhen, China). The images were taken under 20× magnification with resolutions of 3024 × 3024 pixels for smartphone images and 3088 × 2076 pixels for microscope images. To capture images of the mounted wings, the slides were elevated on half a petri dish and illuminated from below using a desktop lamp, as described by Nolte et al (2024). All wing images including meta information have been deposited in a publicly available archive (Nolte et al. 2025) and can be assigned via an unique identifier (image ID) in the supplementary Table S1. For each wing image, landmark coordinates on 18 anatomical junctions of the wing veins (Fig. 1) were digitized with Fiji (Schindelin et al. 2012) as bioscience bundle of ImageJ (Schneider et al. 2012). The chosen landmark configuration is in accordance with other mosquito studies employing geometric morphometrics (Wilke et al. 2016, Sauer et al. 2020, Martinet et al. 2021). The landmark coordinates were collected by a single experienced observer with entomological background (author HJ). In addition, six different observers (authors FGS, KN, LD, PD, PM and T,) collected landmarks from a subsample of 20 images per species and imaging device to assess the degree of observer error (Table S.X).

**Figure 1:**
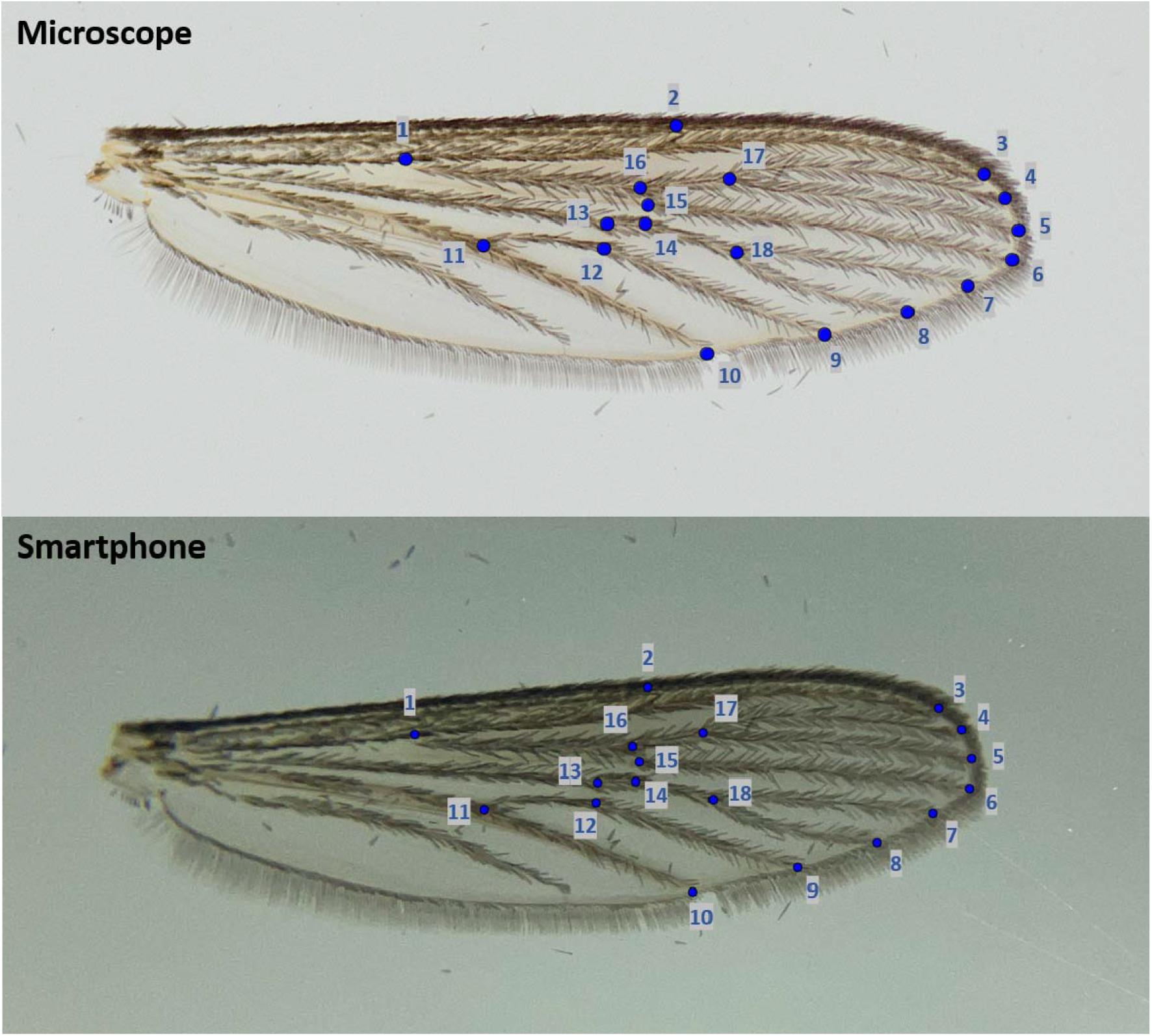
Example wing images captured using a microscope and smartphone, showing the 18 landmarks.

### Statistics

The raw two-dimensional landmark coordinates (Table S1) were superimposed by generalised Procrustes analyses using the R package geomorph (Baken et al. 2021, Adams et al. 2023). The generalised Procrustes analysis was performed separately to the three landmark data sets: microscope images, smartphone images and the combined data set with images from both devices. This procedure was applied independently to landmark datasets generated by a single observer and to those generated by six different observers. During a generalised Procrustes analysis, raw landmark coordinates are centred, scaled and rotated, so that the resulting Procrustes coordinates describe the wing shape in itself (Rohlf and Slice 1990). These shape coordinates were subsequently used to evaluate the species classification accuracy based on images acquired with different imaging devices and observer setups (one and six observers). Four classification algorithms were compared including linear discriminant analysis (LDA) (Venables and Ripley 2002), RF (Liaw and Wiener 2002), supported vector machine (SVM) (Meyer et al. 2019), and XGBoost (Chen et al. 2024). Default hyperparameters were used to ensure comparability between classifiers. For each algorithm, model performance was assessed with an 80:20 training-validation split, which was repeated 20 times using a random data split generated by means of the R package caret, with reproducibility ensured by setting a fixed random seed (Kuhn 2008). The data were split by specimen to avoid that landmark data sets taken by different observer or based on images of the same specimen acquired with two devices, were included in both the training and the validation set. Analyses of variances (ANOVAs) were used to test for statistically significant differences in the obtained species classification accuracies in dependence of the imaging devices (categories: microscope, smartphone or both) and classification algorithm (categories: LDA, RF, SVM, or XGBoost). Separate ANOVAs were conducted for landmark datasets obtained by a single observer and by six observers.

To evaluate the effects of digitization and biological variation on wing shape in the landmark data set obtained by six observers, a Procrustes ANOVA was performed using the procD.lm function in the R package geomorph (Baken et al. 2021, Adams et al. 2023). The model included mosquito species, imaging device, observer, and image ID (entered as a factor to account for the digitization precision for the same wing images) as explanatory variables. Statistical significance was assessed using 999 permutation iterations. Sequential (Type I) sums of squares were used to partition shape variation among factors. Following the approach by Klingenberg and McIntyre (1998), we subsequently calculated the sum of squares and the mean sum of squares separately for each landmark. This can help to localize the landmark-wise shape variation associated with the different factors. However, we are aware that these results should be interpreted with caution as generalized Procrustes analysis reduces the effect of error-prone landmarks with high inaccuracy, and thereby increase the variability among the remaining landmarks (von Cramon□Taubadel et al. 2007). This phenomenon was originally described by Chapman (1990) as the “Pinocchio effect”. Therefore, to further assess measurement errors, we calculated the pixel error for each landmark using the raw non-superimposed landmark coordinates obtained from the multi-observer data set. Specifically, the six measurements by the different observers per image were used to calculate the mean coordinate for each landmark. These were used to compute the Euclidean pixel distance between single landmark measurements and their corresponding mean landmark, serving as a proxy for landmark-wise deviation among observers (Messer et al. 2022). All statistical analyses were conducted in R (R Core Team 2022) and visualized by the R package “ggplot2” (Wickham 2016).

## Results

Species classification accuracy demonstrated substantial variability, ranging from 55% to 100% depending on the landmark data sets and classification algorithms applied (Fig. 2). For the landmark data collected by a single observer, both the chosen imaging device (F_2,234_ = 59.1, p < 0.0001) and the chosen classification algorithm (F_3,234_ = 33.2, p < 0.0001) had significant effects on the classification accuracy. Thereby, the highest accuracy was observed for landmarks collected from microscope images, and among the four classification algorithms, LDA consistently provided higher accuracies (Fig. 2). Misclassification occurred predominantly between *Ae. japonicus* and *Ae. koreicus*, with classification accuracies ranging from 80% for smartphone-image derived landmarks to 92% based on the landmarks from microscope images. *Aedes aegypti* and *Ae. albopictus* were classified with high accuracy ranging from 92% to 99% (Table 1, Table S2).

**Table 1:** Mean confusion matrix obtained through 20-fold cross validation using four different classifiers (LDA: Linear Discriminant Analysis; SVM: Supported Vector Machine; RF: Random Forest; XGB: XGBoost). The matrix is based on the single observer landmark data combining microscope and smartphone images. Green cells indicate correct classification, whereas yellow cells indicate wrong misclassification.

|  | Classifiers | *Ae. aegypti* | *Ae. albopictus* | *Ae. japonicus* | *Ae. koreicus* |
| --- | --- | --- | --- | --- | --- |
| *Ae. aegypti* | LDA | 98.01% | 2.38% | 0.92% | 1.64% |
|  | SVM | 96.10% | 2.54% | 1.46% | 1.89% |
|  | RF | 95.41% | 4.34% | 1.15% | 2.21% |
|  | XGB | 95.68% | 3.52% | 0.92% | 1.48% |
| *Ae. albopictus* | LDA | 0.68% | 97.05% | 0.54% | 0.49% |
|  | SVM | 1.58% | 96.39% | 1.92% | 0.33% |
|  | RF | 2.26% | 94.10% | 1.85% | 0.66% |
|  | XGB | 1.64% | 94.26% | 1.08% | 1.15% |
| *Ae. japonicus* | LDA | 1.16% | 0.16% | 89.15% | 10.00% |
|  | SVM | 1.37% | 0.49% | 86.31% | 8.93% |
|  | RF | 1.16% | 1.56% | 88.31% | 10.98% |
|  | **XGB** | 1.37% | 1.56% | 88.38% | 10.25% |
| ***Ae. koreicus*** | **LDA** | 0.89% | 0.66% | 10.31% | 90.49% |
|  | **SVM** | 0.34% | 1.15% | 12.00% | 88.77% |
|  | **RF** | 1.10% | 0.08% | 8.69% | 87.79% |
|  | **XGB** | 1.23% | 0.74% | 9.62% | 89.51% |

**Figure 2:**
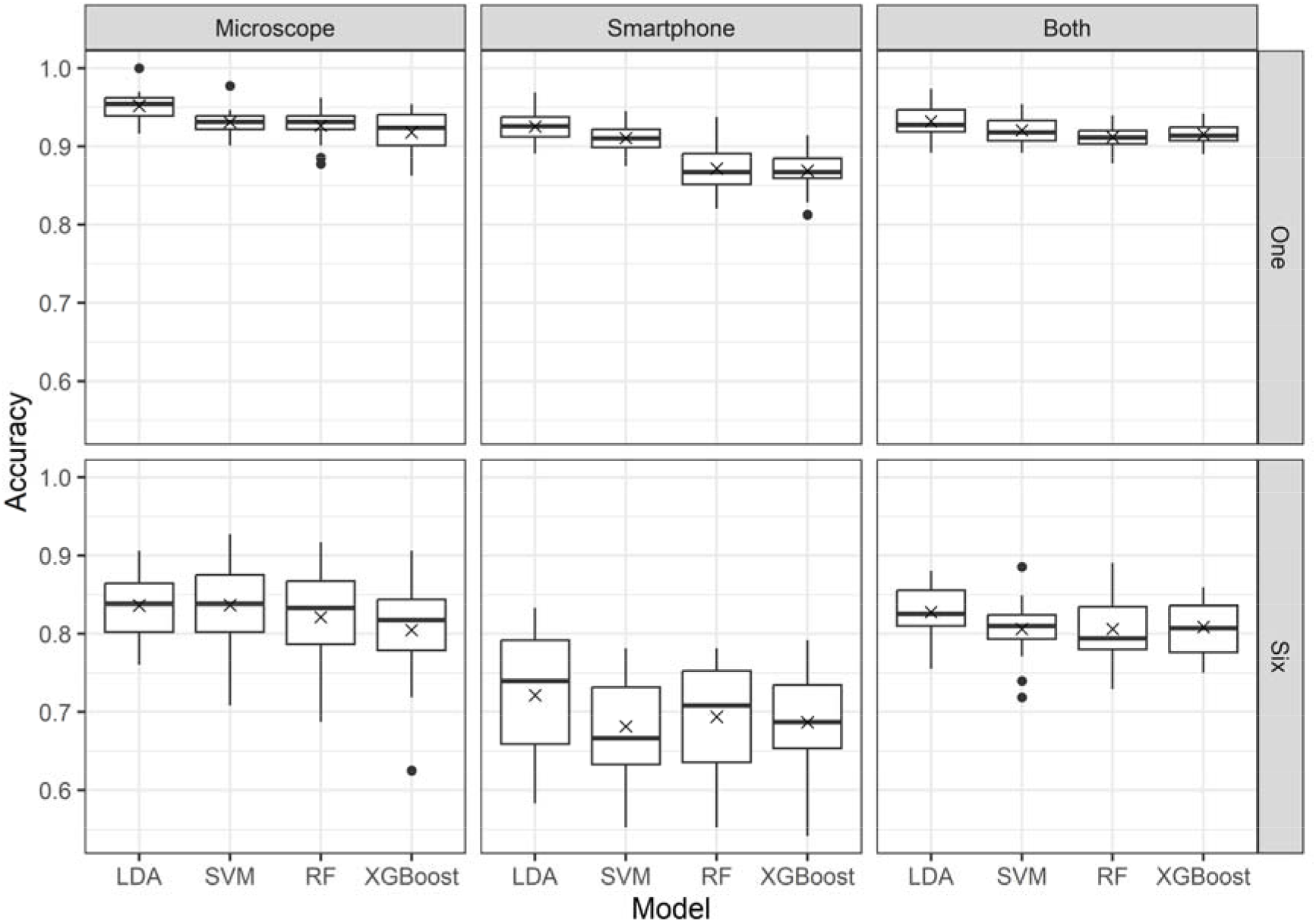
Boxplots showing the species classification accuracy obtained with linear discriminant analysis (LDA), Random Forest (RF), supported vector machine (SVM) and XGBoost. The algorithms were applied to shape coordinates derived from landmarks collected by either a single observer on the full data set (one) or by six observers using a subset of 20 images per species and device (six). Results are shown separately for microscope images (left), smartphone images (center), or a combination of both image types (right). The “X” denotes the mean.

The classification accuracy based on the landmark data collected by six observers was also significantly affected by the imaging device (F_2,234_ = 131.1, p < 0.0001) and classification algorithm (F_3,234_ = 2.9, p = 0.0357) (Fig. 2). In the multi-observer data, LDA achieved slightly higher accuracy for the landmarks derived from smartphone images than the other classifiers. No clear performance differences among classifiers were observed for landmarks derived from microscope images. (Fig. 2).

The Procrustes ANOVA revealed statistically significant effects of species, image ID, device and observer on wing shape variation, accounting for 74.8 % of the total variation (Table 2). Variation among the image ID accounted for the largest proportion (41.7 %), but the mean square was comparatively low (0.01) compared to other factors. This indicates that the total variation attributed to Image ID is distributed across 155 degrees of freedom (Table 2), meaning that individual image effects are small but collectively contribute substantially to the overall variance. Species identity explained a statistically significant and large proportion of the shape variation (24.9 %) (Table 2). Small but also significant effects were detected for the used imaging device (0.5 %) and the observer (7.7 %) (Table 2).

**Table 2:**
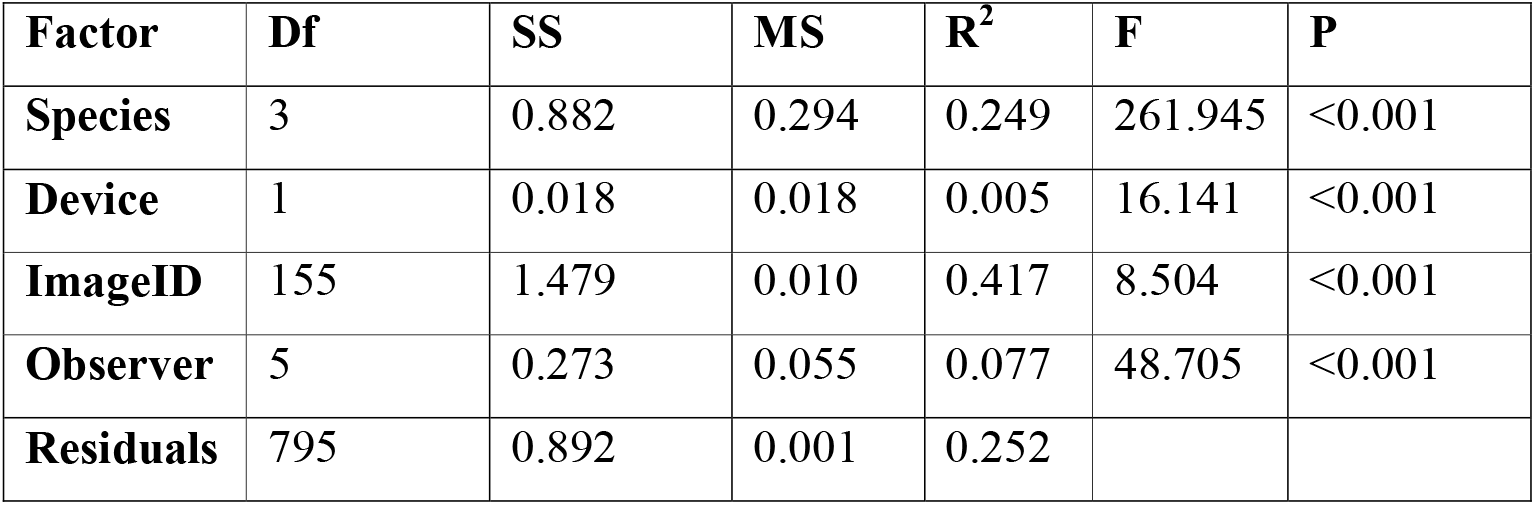
Results of the Procrustes ANOVA including degrees of freedom (Df), sum of squares (SS), mean square (MS), R^2^, F-statistic (F) and p value (P) for each explanatory variable.

| **Factor** | **Df** | **SS** | **MS** | **$R^2$** | **F** | **P** |
| --- | --- | --- | --- | --- | --- | --- |
| **Species** | 3 | 0.882 | 0.294 | 0.249 | 261.945 | <0.001 |
| **Device** | 1 | 0.018 | 0.018 | 0.005 | 16.141 | <0.001 |
| **ImageID** | 155 | 1.479 | 0.010 | 0.417 | 8.504 | <0.001 |
| **Observer** | 5 | 0.273 | 0.055 | 0.077 | 48.705 | <0.001 |
| **Residuals** | 795 | 0.892 | 0.001 | 0.252 |  |  |

The mean squares per landmark varied substantially among the different factors and landmarks (Fig. 3). Landmark 2, 17 and 18 exhibited the greatest variability among species (Fig. 3, Table S3). Landmark 13 showed the strongest variability associated with the observer factor, potentially indicating systematic differences in landmark placement among observers (Fig. 3, Table S3). In addition, landmark 2 displayed the highest variability associated with the image ID, suggesting considerable variation among the repeated measurements of the same image (Fig. 3, Table S3). This observation is also reflected in the Euclidean pixel distances calculated based on the raw landmark coordinates, where the highest observer-related variation was detected at landmark 2 (Fig. 4). Moreover, this analysis demonstrated that smartphone images consistently yielded greater inter-observer pixel distances than microscope images across all landmarks (Fig. 4).

**Figure 3:**
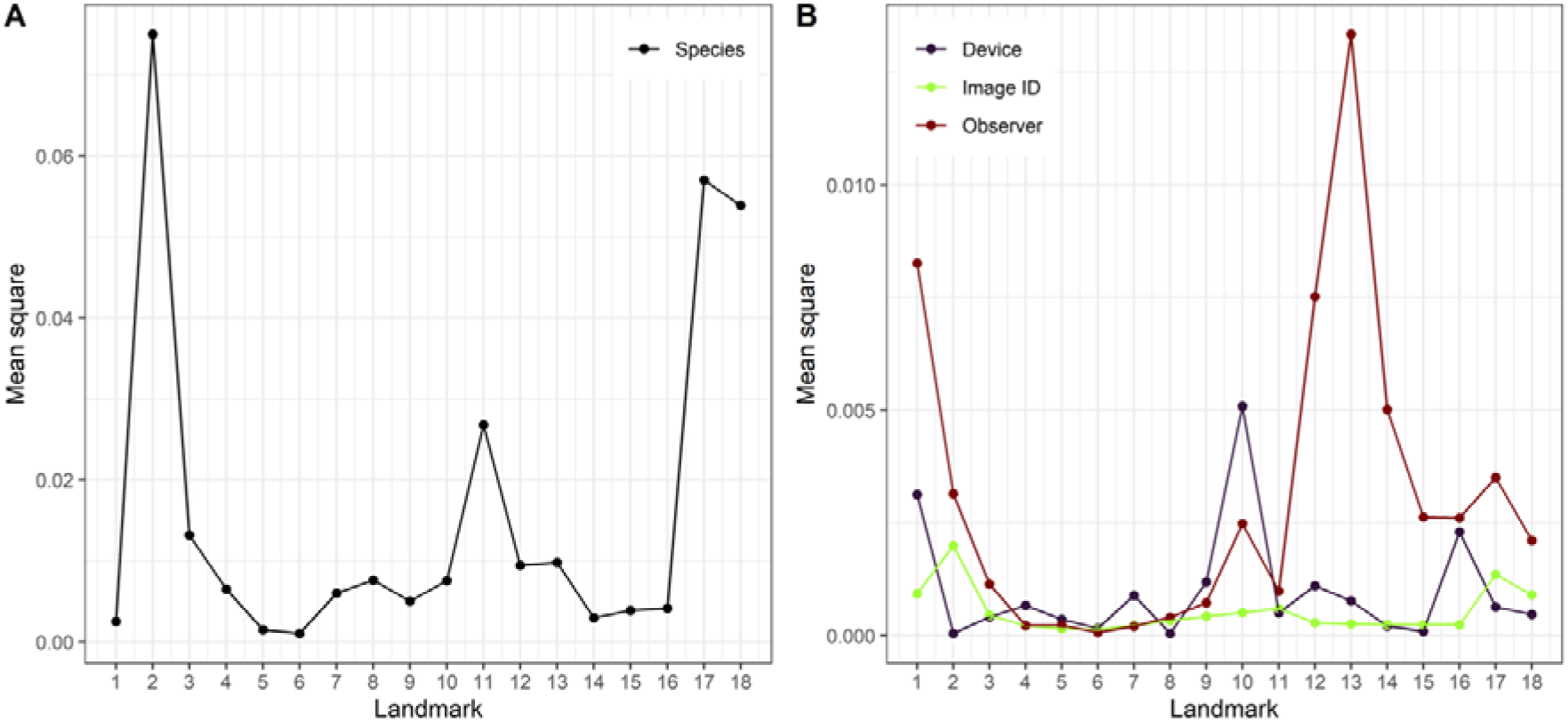
Mean square values representing the amount of variation for 18 landmarks for the effects of species, device, image ID and observer. Due to differences in magnitude, the effects of species (A) and the effects of device, image ID and observer (B) are presented separately for clarity.

**Figure 4:**
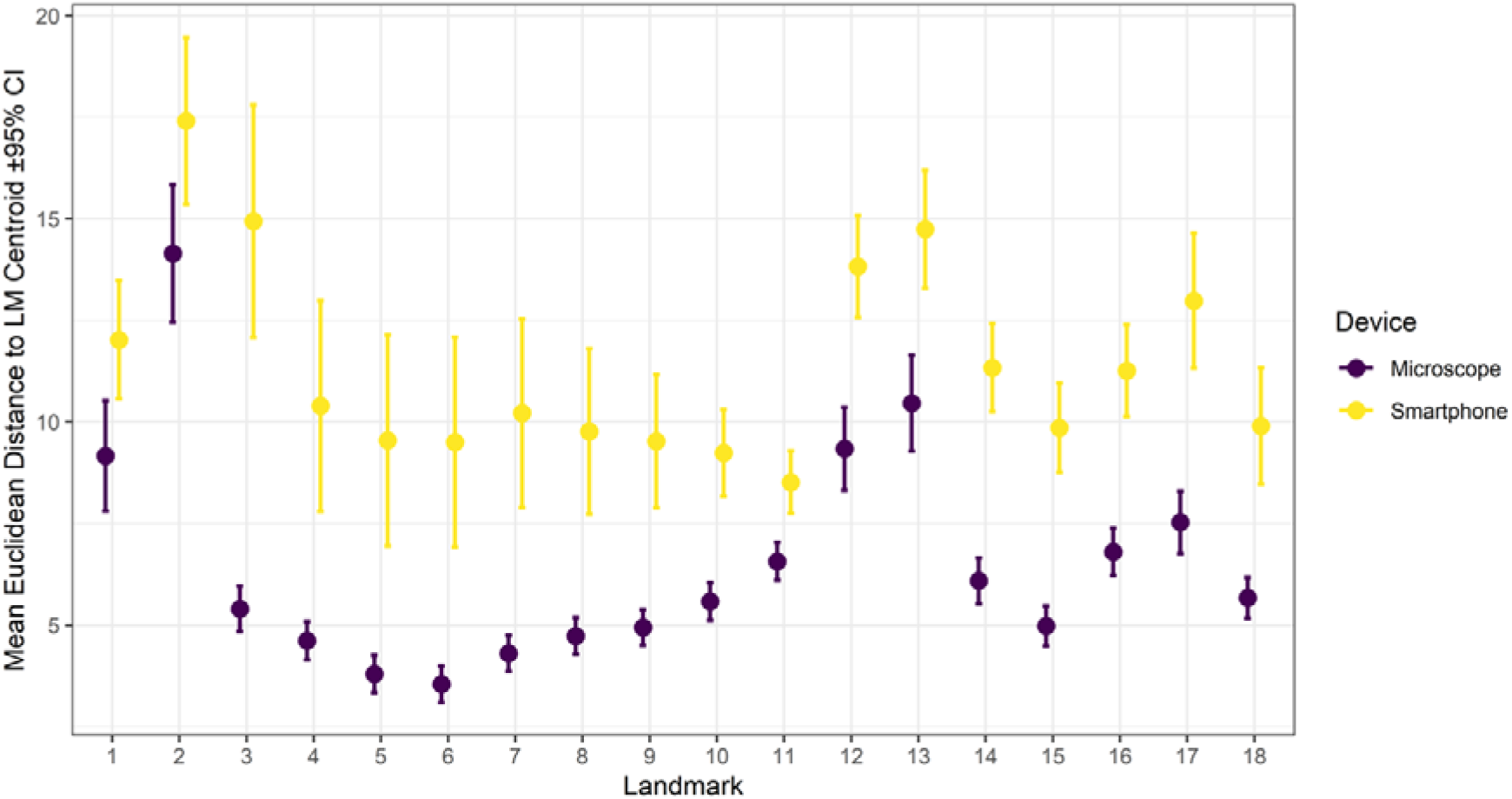
Mean Euclidean pixel distance to the centroid with 95% confidence interval (CI) for each of the 18 landmarks (LM) given for microscope and smartphone images. The figure is based on the multi-observer data, in which the landmarks of 160 images were collected by six different observers.

## Discussion

In line with previous studies (Henry et al. 2010, Sauer et al. 2023), the four *Aedes* species could be reliably distinguished based on the analysis of their wing shape, with slightly lower accuracy for *Ae. japonicus* and *Ae. koreicus* (>90%) compared to *Ae. aegypti* and *Ae. albopictus* (>95%). However, our analyses demonstrated notable variation in the species classification accuracies in dependence of the choice of the imaging device, the number of observers and the classification algorithm.

In the data set generated by a single observer, the LDA achieved the highest accuracy among the four evaluated classification algorithms. This result is consistent with a geometric morphometric study on the cockle *Cerastoderma edule*, in which LDA likewise outperformed the other twelve classification algorithms used to distinguish different populations based on their shell shape (Martins et al. 2024). Procrustes coordinates often approximate multivariate normality with comparable covariance structures among all classes, closely matching the assumptions of LDA. Under such conditions, simpler parametric methods can outperform more complex, nonlinear classification algorithms such as RF (Guo et al. 2010). In the multi-observer dataset, the effect of the chosen classifier was less pronounced and RF, SVM, and XGBoost performed similar to LDA. Small discrepancies in the landmark placement due to different observers introduce skewness in the data, potentially resulting in deviations from multivariate normality. Machine learning classifiers such as RF are robust to non-normal distributed data (Guo et al. 2010, Khondoker et al. 2016, Chowdhury et al. 2022), and are therefore likely to be less sensitive to the observer-induced variability in the landmark data.

In geometric morphometrics, observer induced errors in landmark collection represents an important source of variation and can substantially limit its broader applicability (von Cramon□Taubadel et al. 2007, Fruciano 2016). A high variation among observers, as indicated by the mean square, was detected at landmark 13. At this landmark, a cross vein connects the cubitus 1 to the media. In the analysed *Aedes* species, this cross vein is very pale and difficult to see, which likely explains why different observers may have interpreted this intersection differently. Landmark 2, the intersection of the costa and the subcosta, showed the highest mean square associated with the image ID. Likewise, the largest mean Euclidean pixel distance among the six observers, calculated from the raw landmark coordinates, was observed at this landmark. This pattern is consistent with a landmark-wise comparison of the observer error in *Anopheles* wings reported by Lorenz and Suesdek (2013). At landmark 2, the subcostal vein gradually merges into the costa, making the exact intersection difficult to determine. In addition, dense scales cover the veins, further complicating the precise identification of this landmark. Lorenz and Suesdek (2013) demonstrated that the chemical or physical removal of wing scales reduces the observer effect. However, for our study, we did not remove the scales to shorten preparation time and to maintain a workflow that allows rapid species classification. Under these conditions, landmark 2 may introduce a relevant Pinocchio effect, i.e. Procrustes analysis reduces the influence of error-prone, highly inaccurate landmarks, thereby increasing the relative variability among the remaining landmarks (von Cramon□Taubadel et al. 2007). Nevertheless, relative to the species-specific variation, the observer effect can still be considered small. The Procrustes ANOVA showed that the mean square associated with methodological factors (repeated measurements, observers or imaging device) was distinctly lower than the mean square among species.

The landmark-wise partitioning of variation revealed that landmark 2, 17, and 18 were most variable among the four *Aedes* species. Previous work also reported a tendency for greater differences at landmark 18 between *Ae. japonicus* and *Ae. koreicus* (Sauer et al. 2023). However, none of the landmarks alone would be sufficient to reliably discriminate these *Aedes* species. Instead, the full set of landmarks is required for species differentiation.

The lower image quality of smartphone images decreases the reproducibility of the landmark collection. However, when the landmarks were collected by a single observer, classification accuracy (∼ 92°%) was only slightly reduced compared to the microscope images (∼ 95°%). This suggests that smartphone-based imaging can provide a highly cost-effective solution for geometric morphometric analyses and may facilitate species identification in low-resource settings. This method could be faster if the wings are not mounted as in our study. Zulzahrin et al. (2024) covered the dissected mosquito wings with immersion oil and a glass cover slip for direct image collection. The authors used the images to measure the wing size, but this approach might also be promising for landmark-based geometric morphometrics and could further accelerate the species identification process, thereby supporting its integration into mosquito surveillance and monitoring programs.

In conclusion, the four most relevant invasive *Aedes* species in Europe can be reliably distinguished by means of geometric morphometrics. The use of a smartphone with a macro lens provided sufficient image quality for accurate landmark collection. However, observer bias was notably higher compared to microscope images. Therefore, when using this low-cost alternative, landmarks should be collected by a single experienced observer.

## Author contribution

FGS, KN, and RL wrote the first manuscript draft. FGS and KN analysed the data. All authors were involved in data collection and revised the manuscript.

