## Supplementary material for "Accurate identification of invasive *Aedes* mosquito species using low-cost imaging and geometric wing morphometrics": Table S2 and Table S3

Tab. S2: mean classification accuracy after 20 validation runs for the landmarks derived from microscope (left) and smartphone (right) images obtained from four different classification algorithms (LDA: Linear Discriminant Analysis, SVM: Supported Vector Machine, RF: Random Forest).

| **LDA: Microscope images** | | | | |
|---|---|---|---|---|
| | *Ae.aegypti* | *Ae.albopictus* | *Ae.japonicus* | *Ae.koreicus* |
| *Ae.aegypti* | 0.997 | 0.013 | 0.005 | 0.016 |
| *Ae.albopictus* | 0.000 | 0.981 | 0.003 | 0.006 |
| *Ae.japonicus* | 0.000 | 0.002 | 0.919 | 0.072 |
| *Ae.koreicus* | 0.003 | 0.005 | 0.073 | 0.906 |
| **balanced accuracy** | ***0.951*** | | | |

| **LDA: Smartphone images** | | | | |
|---|---|---|---|---|
| | *Ae.aegypti* | *Ae.albopictus* | *Ae.japonicus* | *Ae.koreicus* |
| *Ae.aegypti* | 0.974 | 0.045 | 0.003 | 0.002 |
| *Ae.albopictus* | 0.019 | 0.943 | 0.009 | 0.000 |
| *Ae.japonicus* | 0.007 | 0.003 | 0.888 | 0.109 |
| *Ae.koreicus* | 0.000 | 0.008 | 0.100 | 0.890 |
| **balanced accuracy** | **0.924** | | | |

| **SVM: Microscope images** | | | | |
|---|---|---|---|---|
| | *Ae.aegypti* | *Ae.albopictus* | *Ae.japonicus* | *Ae.koreicus* |
| *Ae.aegypti* | 0.979 | 0.015 | 0.016 | 0.022 |
| *Ae.albopictus* | 0.008 | 0.960 | 0.022 | 0.005 |
| *Ae.japonicus* | 0.004 | 0.010 | 0.883 | 0.077 |
| *Ae.koreicus* | 0.008 | 0.016 | 0.080 | 0.897 |
| **balanced accuracy** | **0.930** | | | |

| **SVM: Smartphone images** | | | | |
|---|---|---|---|---|
| | *Ae.aegypti* | *Ae.albopictus* | *Ae.japonicus* | *Ae.koreicus* |
| *Ae.aegypti* | 0.953 | 0.048 | 0.011 | 0.010 |
| *Ae.albopictus* | 0.033 | 0.945 | 0.018 | 0.000 |
| *Ae.japonicus* | 0.013 | 0.000 | 0.856 | 0.105 |
| *Ae.koreicus* | 0.001 | 0.007 | 0.115 | 0.884 |
| **balanced accuracy** | **0.910** | | | |

| **RF: Microscope images** | | | | |
|---|---|---|---|---|
| | *Ae.aegypti* | *Ae.albopictus* | *Ae.japonicus* | *Ae.koreicus* |
| *Ae.aegypti* | 0.961 | 0.016 | 0.013 | 0.020 |
| *Ae.albopictus* | 0.018 | 0.971 | 0.011 | 0.006 |
| *Ae.japonicus* | 0.010 | 0.013 | 0.909 | 0.113 |
| *Ae.koreicus* | 0.011 | 0.000 | 0.067 | 0.861 |
| **balanced accuracy** | **0.926** | | | |

| **RF: Smartphone images** | | | | |
|---|---|---|---|---|
| | *Ae.aegypti* | *Ae.albopictus* | *Ae.japonicus* | *Ae.koreicus* |
| *Ae.aegypti* | 0.928 | 0.070 | 0.033 | 0.010 |
| *Ae.albopictus* | 0.039 | 0.922 | 0.024 | 0.002 |
| *Ae.japonicus* | 0.031 | 0.005 | 0.789 | 0.143 |
| *Ae.koreicus* | 0.003 | 0.003 | 0.153 | 0.845 |
| **balanced accuracy** | **0.871** | | | |

| **XGBoost: Microscope images** | | | | |
|---|---|---|---|---|
| | *Ae.aegypti* | *Ae.albopictus* | *Ae.japonicus* | *Ae.koreicus* |
| *Ae.aegypti* | 0.9639 | 0.0210 | 0.0188 | 0.0219 |
| *Ae.albopictus* | 0.0153 | 0.9694 | 0.0109 | 0.0047 |
| *Ae.japonicus* | 0.0056 | 0.0048 | 0.8781 | 0.1172 |
| *Ae.koreicus* | 0.0153 | 0.0048 | 0.0922 | 0.8563 |
| **balanced accuracy** | **0.9169** | | | |

| **XGBoost: Smartphone images** | | | | |
|---|---|---|---|---|
| | *Ae.aegypti* | *Ae.albopictus* | *Ae.japonicus* | *Ae.koreicus* |
| *Ae.aegypti* | 0.924 | 0.067 | 0.036 | 0.010 |
| *Ae.albopictus* | 0.032 | 0.918 | 0.021 | 0.012 |
| *Ae.japonicus* | 0.031 | 0.005 | 0.792 | 0.140 |
| *Ae.koreicus* | 0.014 | 0.010 | 0.150 | 0.838 |
| **balanced accuracy** | **0.868** | | | |

Table S3: Sum of squares (SS) and mean squares (MS) for the effects of the Procrustes ANOVA (Table 2), calculated separately for each of the 18 landmarks. Increasing red color intensity indicates higher SS and MS values.

| Landmark | SS Device | SS species | SS Image | SS Observer | SS error | MS Device | MS species | MS Image | MS Observer | MS error |
|---|---|---|---|---|---|---|---|---|---|---|
| 1 | 0.00313 | 0.00756 | 0.14405 | 0.04132 | 0.08514 | 0.00313 | 0.00252 | 0.00093 | 0.00826 | 0.00011 |
| 2 | 0.00005 | 0.2251 | 0.30966 | 0.01574 | 0.23063 | 0.00005 | 0.07503 | 0.002 | 0.00315 | 0.00029 |
| 3 | 0.00041 | 0.03951 | 0.07095 | 0.00572 | 0.07463 | 0.00041 | 0.01317 | 0.00046 | 0.00114 | 0.00009 |
| 4 | 0.00067 | 0.01954 | 0.03419 | 0.00112 | 0.01426 | 0.00067 | 0.00651 | 0.00022 | 0.00022 | 0.00002 |
| 5 | 0.00036 | 0.0045 | 0.0242 | 0.00117 | 0.01049 | 0.00036 | 0.0015 | 0.00016 | 0.00023 | 0.00001 |
| 6 | 0.00016 | 0.00308 | 0.02097 | 0.00035 | 0.01134 | 0.00016 | 0.00103 | 0.00014 | 0.00007 | 0.00001 |
| 7 | 0.0009 | 0.01804 | 0.03464 | 0.00104 | 0.01472 | 0.0009 | 0.00601 | 0.00022 | 0.00021 | 0.00002 |
| 8 | 0.00005 | 0.02284 | 0.05172 | 0.00204 | 0.01729 | 0.00005 | 0.00761 | 0.00033 | 0.00041 | 0.00002 |
| 9 | 0.00119 | 0.01512 | 0.06681 | 0.00367 | 0.02252 | 0.00119 | 0.00504 | 0.00043 | 0.00073 | 0.00003 |
| 10 | 0.00509 | 0.02266 | 0.07928 | 0.01242 | 0.0291 | 0.00509 | 0.00755 | 0.00051 | 0.00248 | 0.00004 |
| 11 | 0.00051 | 0.08042 | 0.09302 | 0.00495 | 0.03695 | 0.00051 | 0.02681 | 0.0006 | 0.00099 | 0.00005 |
| 12 | 0.00111 | 0.02837 | 0.04407 | 0.03763 | 0.06237 | 0.00111 | 0.00946 | 0.00028 | 0.00753 | 0.00008 |
| 13 | 0.00077 | 0.02945 | 0.04064 | 0.06672 | 0.07543 | 0.00077 | 0.00982 | 0.00026 | 0.01334 | 0.00009 |
| 14 | 0.00021 | 0.00892 | 0.03857 | 0.0251 | 0.03207 | 0.00021 | 0.00297 | 0.00025 | 0.00502 | 0.00004 |
| 15 | 0.0001 | 0.01164 | 0.03814 | 0.01317 | 0.03065 | 0.0001 | 0.00388 | 0.00025 | 0.00263 | 0.00004 |
| 16 | 0.0023 | 0.01246 | 0.03793 | 0.01306 | 0.04517 | 0.0023 | 0.00415 | 0.00024 | 0.00261 | 0.00006 |
| 17 | 0.00064 | 0.17109 | 0.21004 | 0.01753 | 0.06461 | 0.00064 | 0.05703 | 0.00136 | 0.00351 | 0.00008 |
| 18 | 0.00047 | 0.16165 | 0.1404 | 0.01057 | 0.03484 | 0.00047 | 0.05388 | 0.00091 | 0.00211 | 0.00004 |
